# Predicting vaccine effectiveness in livestock populations: a theoretical framework applied to PRRS virus infections in pigs

**DOI:** 10.1101/563205

**Authors:** Vasiliki Bitsouni, Samantha Lycett, Tanja Opriessnig, Andrea Doeschl-Wilson

## Abstract

Vaccines remain one of the main tools to control infectious diseases in domestic livestock. Although a plethora of veterinary vaccines are on the market and routinely applied to protect animals against infection with particular pathogens, the disease in question often continues to persist, sometimes at high prevalence. The limited effectiveness of certain vaccines in the field leaves open questions regarding the required properties that an effective vaccine should have, as well as the most efficient vaccination strategy for achieving the intended goal of vaccination programmes. To date a systematic approach for studying the combined effects of different types of vaccines and vaccination strategies is lacking. In this paper, we develop a theoretical framework for modelling the epidemiological consequences of vaccination with imperfect vaccines of various types, administered using different strategies to herds with different replacement rates and heterogeneity in vaccine responsiveness. Applying the model to the Porcine Reproductive and Respiratory Syndrome (PRRS), which despite routine vaccination remains one of the most significant endemic swine diseases worldwide, we then examine the influence of these diverse factors alone and in combination, on within-herd virus transmission. We derive threshold conditions for preventing infection invasion in the case of imperfect vaccines inducing limited sterilizing immunity. The model developed in this study has practical implications for the development of vaccines and vaccination programmes in livestock populations not only for PRRS, but also for other viral infections.

## Introduction

For decades, vaccination has been considered the most powerful defense against a range of infectious diseases. The major aims of veterinary vaccines are to improve the health of animals and to prevent or reduce pathogen transmission, thereby mitigating the impact of infectious disease on livestock production in a cost-effective manner [1]. However, the potential of available vaccines to effectively control infectious disease in livestock is contentious [2], because they often only confer limited sterilizing immunity and thus may not prevent infection, and may only partly reduce pathogen transmission. A vaccine is considered effective if it can reduce within-host pathogen burden as well as pathogen shedding, prevent or alleviate disease-induced clinical signs, and thus improve the general health conditions of exposed animals [1]. More comprehensively, the desirable properties of an effective vaccine include: (i) high safety (i.e. no reversion to virulence or disease caused by the vaccine strain) [3–5]; (ii) high sterilizing immunity against a wide range of variant pathogen strains [6,7]; (iii) fast onset of protection [8]; (iv) high immunogenicity leading to reduction in pathogen load, shedding and faster recovery [9], as well as (v) vaccine responsiveness in a broad range of hosts. Very few vaccines on the market satisfy all of these properties. For example, vaccine safety is a major concern for modified live vaccines [1,10,11], sterilizing immunity has been found to often reach alarmingly low values [12], and heterogeneity in vaccine response, e.g. due to genetic or age differences, seems ubiquitous [13–15].

All of these listed vaccine properties play an important role in pathogen transmission, and thus in vaccine effectiveness on a population level. For example, sterilizing immunity affects the susceptibility of a host to infection with a heterologous strain, whereas the impact of a vaccine on pathogen shedding affects an individual’s infectivity, i.e. its ability to transmit infection to others [16]. Vaccines that accelerate host recovery reduce pathogen transmission by reducing the infectious period of a host [16–18]. In contrast, delay in onset of protection or host heterogeneity in vaccine response limit the time or extent of effective vaccine coverage in a population, thus enabling continued pathogen transmission.

The effectiveness of a given vaccine in the field depends not only on the properties of the vaccine itself, but also on how the vaccine is applied and what other biosecurity measures are in place. For example, herd closure during a disease outbreak has been promoted as a highly effective disease control strategy, whereas continuous influx of new susceptible, possibly non-vaccinated individuals contributes to long term persistence of the disease in a herd [17,19]. Common vaccination strategies for livestock diseases include prophylactic (also known as preventative) and reactive mass vaccination [20]. Prophylactic vaccination is applied prior to introduction of a pathogen into a herd, typically either as precaution to avoid recurrence of previously resolved disease outbreaks in the herd or due to a perceived high risk occurring from outbreaks in neighboring herds or farms. Although prophylactic mass vaccination is rare in practice, it is considered the best strategy to prevent disease outbreaks and thus to minimize the risk of a major epidemic [2]. Reactive vaccination on the other hand, although considered less effective than prophylactic vaccination, is typically applied to control ongoing epidemics. Application of either vaccination strategy is commonly hampered in practice by insufficient vaccine availability, economic reasons or safety restrictions (e.g. clinical symptoms occurring due to the vaccination) and logistic delays [20]. These affect the frequency and timing of vaccination as well as the effective vaccine coverage in a population, i.e. the proportion of immunized animals. One-off vaccination of a small proportion of animals in a herd with high disease prevalence and high replacement rate may not be very effective, even if the applied vaccine contains all the desirable properties.

To the best of our knowledge, there is currently no theoretical framework to systematically assess how different vaccine properties and vaccination strategies interactively influence infection invasion and transmission in herds with different demographics. This makes predictions of vaccine effectiveness in the field extremely difficult. In particular, an infectious disease for which a comprehensive framework that combines the diverse factors compromising vaccine effectiveness would be extremely useful, is the Porcine Reproductive and Respiratory Syndrome (PRRS), which is caused by the PRRS virus (PRRSv) [21,22]. PRRS is one of the most significant and costly swine diseases globally [23], with estimated costs per year over $ 650 million in the United states [24] and almost 1.5b € in Europe [25]. Although the first PRRS vaccine has become commercially available more than two decades ago and several PRRS vaccines have been widely used since, the prevalence of PRRS remains high [26]. Failed vaccination programmes have raised an urgent demand for more effective vaccines [27,28]. As PRRS continues to spread rapidly all over the world, with more virulent PRRSv strains emerging in Asia a few years ago, concerns have been raised about the epidemiological consequences of vaccination and the evaluation of the vaccination effectiveness [29,30].

The most common PRRS vaccines can be broadly categorised into modified-live virus (MLV) vaccines or inactivated or killed virus (KV) vaccines. However, so far none of them has been fully effective in preventing the spread of the virus within a herd [8,21,22,26,31,32]. PRRS MLV vaccines are attenuated live vaccines, which have shown delayed but effective protection against homologous and some heterologous PRRSv strains [8]. They reduce clinical signs, severity and the duration of viraemia and virus shedding [31, 33]. However, their limited sterilizing immunity and immunogenicity to many circulating PRRSv strains have raised major concerns regarding vaccine effectiveness [8,21,34–37]. PRRS KV vaccines, containing adjuvants, on the other hand are known for their high safety, but the sterilizing immunity that they provide against either homologous or heterologous PRRSv strains is extremely limited, and they often fail to significantly reduce clinical signs [38], viremia and duration of shedding in nave animals [39]. While KV vaccines have failed to elicit detectable antibodies and also barely elicit cell-mediated immune response (CMI) response in PRRSv-negative pigs (i.e. ineffective prophylactic vaccination) [40], in PRRSv-positive pigs (reactive vaccination) they have been reported to strengthen both types of immune responses to the infecting virus [40,41], thus speeding up of recovery and potentially also reducing infectivity of the pigs. For this reason, PRRS KV vaccines have been recommended for use as therapeutic vaccines for PRRSv treatment rather than for disease prevention [42].

Despite tremendous efforts over the last three decades to understand PRRS pathogenesis and vaccinology, effective PRRS vaccines, possessing safety, broad sterilizing immunity and high immunogenicity, are still lacking [26,28]. Furthermore, relatively little is known how the existing vaccines affect virus transmission in a herd, or how these could be most effectively applied to prevent PRRS outbreaks or reduce their impact. Experimental or field studies testing the impact of a vaccine on virus transmission are not only rare but are limited to a specific vaccine type, a specific vaccination strategy, a specific challenge strain, and specific pig breeds [43]. Mathematical models, on the other hand, have been proven powerful tools to assess the combined effects of several interacting factors on virus transmission and to predict the outcome of different types of vaccines or vaccination strategies (see e.g. [2,44,45] and other references therein).

The aim of this study was to develop a theoretical framework for modelling the combined epidemiological consequences of different vaccination strategies and different vaccine properties applied to domestic livestock populations with different replacement rates. Table 1 lists the key factors known to compromise vaccine effectiveness that are considered in this study. Vaccine safety is not included in this table as in this study we only consider safe vaccines, i.e. vaccines that do not revert to virulence and cause disease by themselves.

**Table 1.** A list of factors that compromise vaccine effectiveness incorporated in our model. Vaccine responsiveness and administration are both captured by the same parameter, *p*, which is the vaccine effective coverage. An incomplete coverage is either due to the vaccine only being administrated to a proportion of pigs or because not all vaccinated pigs have developed protective immunity. A detailed description of the model parameters is given in Table 2.

| Factors | Effect on epidemiological characteristics | Model parameters |
| --- | --- | --- |
| **Vaccine properties** |  |  |
| Vaccine-induced sterilizing immunity | Reduced transmission through lower host susceptibility | $\epsilon_s$ |
| Vaccine-induced reduction in pathogen shedding | Reduced transmission through lower host infectivity | $\epsilon_i$ |
| Vaccine-induced increase in recovery rate | Speed up recovery rate of infected host | $\epsilon_\gamma$ |
| Time delay in onset of immunity | Period between vaccination and the onset of vaccine-induced immunity | time-dependent $\epsilon_{s,i,\gamma}(t)$ |
| Effective coverage (responsiveness) | Proportion of immunized pigs | $p$ |
| **Vaccination strategies** |  |  |
| Prophylactic/Reactive * | Vaccination before/after first infection incident | $\epsilon_{s,i,\gamma}$ |
| Time & frequency | Continuous vaccination/One-off vaccination | $\lambda_N, \lambda_V$ |
| Effective coverage (administration) | Proportion of pigs to which the vaccine is administered | $p$ |
| **Pathogen biology** |  |  |
| Pathogen virulence | Pathogen controlled infection transmission potential | $R_0$ |
| **Herd demography & management** |  |  |
| Replacement rate | Rate at which vaccinated/non-vaccinated pigs are replaced (zero in closed herds) | $\lambda_N, \lambda_V, \mu$ |

\* Note that prophylactic versus reactive vaccination strategies are mainly described by the initial conditions (ICs) of the system as given in the following section.

The core model developed in this study is generic to represent infection dynamics in different livestock populations and for different pathogen species. However, to investigate the epidemiological consequences of vaccination, we parameterize the model to represent a herd of pigs exposed to a PRRSv strain different to the vaccine strain, when vaccination utilizing different types of vaccines (outlined in Table 1) is applied either prophylactically or reactively, at different time points and different frequencies, in herds with different replacement rates. In particular, we use the model to derive threshold conditions for preventing infection invasion even for vaccines with low sterilizing immunity. The model derived in this study provides new insights for vaccine development and application for combating PRRS and other diseases threatening domestic livestock populations.

## Materials and Methods

### Modelling transmission dynamics in a vaccinated population

The generic deterministic epidemiological Susceptible-Infected-Recovered (SIR) model presented below models transmission of a wild-type strain of a particular pathogen in a commercial herd where vaccination is applied.

For an epidemiological SIR model in a homogenous non-vaccinated herd, the transmission dynamics is controlled by two model parameters, the transmission term *β* and the recovery rate *γ*. The transmission term *β*, loosely called transmission rate, is defined as the product of the average contact rate and the probability that virus transmission occurs between an infected and a susceptible hosts upon contact. In this study a frequency-dependent transmission, where the number of contacts is independent of population size, is assumed.

Vaccination introduces heterogeneity in a population, dividing the proportions of pigs into susceptible non-vaccinated (*S_N_*), susceptible vaccinated (*S_V_*), infected non-vaccinated (*I_N_*), infected vaccinated (*I_V_*), and recovered non-vaccinated (*R_N_*) and recovered vaccinated (*R_V_*). All recovered hosts (vaccinated or not) are assumed to gain full immunity to the modelled pathogen field strain. The model is illustrated in Fig 1 and represented by the following set of ordinary differential equations

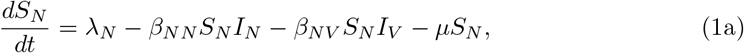

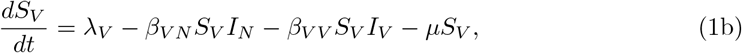

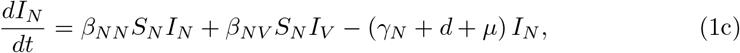

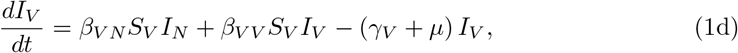

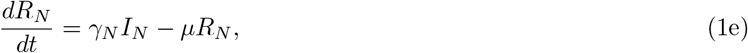

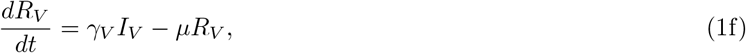

with the initial conditions (ICs):

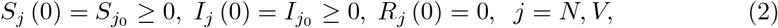

where *β_jk_,j,k* = *N, V*, are the transmission rates from an infected pig of type *k* to a susceptible pig of type *j, γ_j_* are the recovery rates, λ_*j*_ are the corresponding birth/replacement rates and *μ* is the average death/removal rate. Moreover, *d* is the mortality rate due to the infection, which is assumed to be non-zero only for non-vaccinated infected pigs. Equal birth/replacement and death/removal rates are assumed, which corresponds to a constant population size when the death rate due to the infection is zero, i.e. *λ_*N*_* + λ_*V*_ = *μ*. The values *S*_*j*_0__ and *I*_*j*_0__ are non-negative real values denoting the initial proportion of susceptible and infected in population. Table 2 presents a list of all model parameters together with their assumed values for the simulations in this study and corresponding information source. For the simulations it is assumed that the infection with the field pathogen strain in a herd is introduced by a proportion of non-vaccinated, *I*_*N*0_, and/or vaccinated, *I*_*V*0_, individuals in an otherwise fully susceptible population. In prophylactic vaccination *I*_*N*_0__ and *I*_*V*0_ can take values greater than or equal to zero (see Fig 4 in Sec. 1 of Results, where *I*_*N*0_ =0 and *I*_*V*_0__ = 0.001 for the case of full vaccine coverage mass vaccination), while in reactive vaccination it is always *I*_*V*_0__ = 0 (see Fig 6 in Sec. 2 of Results, where *I*_*N*_0__ = 0.001 and *I*_*V*_0__ = 0).

**Fig 1.**
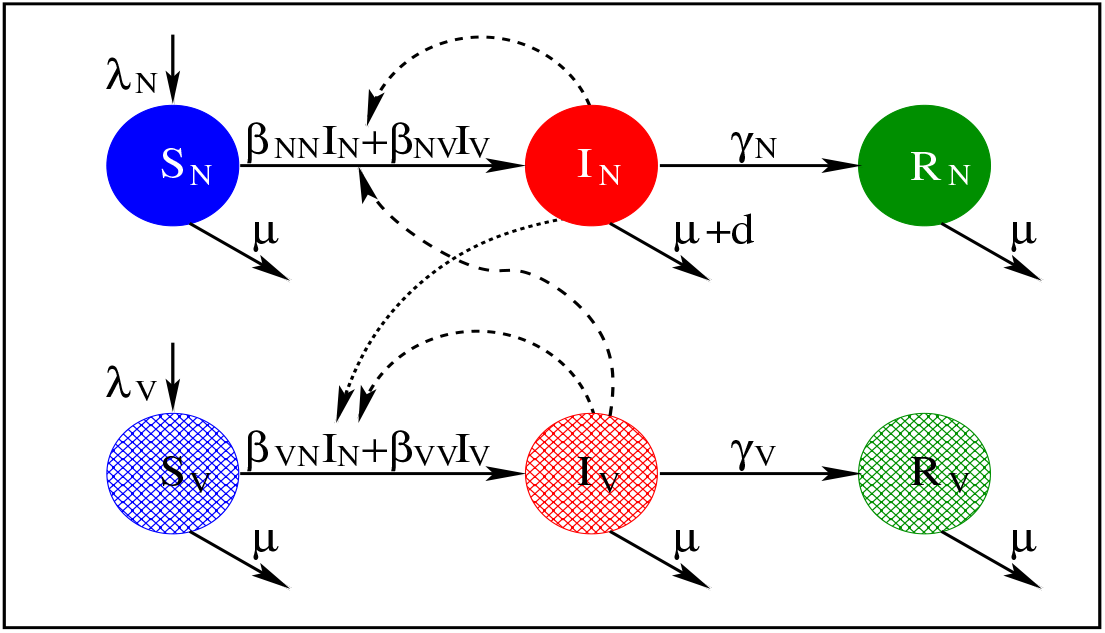
Flow diagrams of the heterogeneous vaccine SIR model given by (1).

**Table 2.** Description of the model parameters together with their assumed value ranges considered in this study.

| Param. | Description | Value | Reference |
| --- | --- | --- | --- |
| $\lambda_N$ | Replacment rate of non-vaccinated susceptible pigs | 0 – 0.0017 | [48–50] |
| $\lambda_V$ | Replacment rate of vaccinated susceptible pigs | 0 – 0.0017 | [48–50] |
| $\beta_{NN}$ | Transmission rate between non-vaccinated pigs | 0.030 – 0.426 | [43, 49, 51, 52] |
| $\beta_{NV}, \beta_{VN}$ | Transmission rates between vac. and non-vac. pigs | 0 – 0.426 | Estimated by relations (7a) & (8a) |
| $\beta_{VV}$ | Transmission rate between vaccinated pigs | 0 – 0.426 | Estimated by relations (7b) & (8b)<br>(see also Ref. [43, 52]) |
| $\gamma_N$ | Recovery rate of non-vaccinated pigs | 0.004 – 0.1428 | [43, 49, 50] |
| $\gamma_V$ | Recovery rate of vaccinated pigs | 0.0057 – 0.1 | Estimated by relation (9)<br>(see also Ref. [43, 49, 53, 54]) |
| $\mu$ | Death/removal rate | 0 – 0.037 | [43, 49, 51, 52] |
| $d$ | Death rate due to infection | 0 – 0.001 | Estimated |
| $R_0^N$ | Basic reproductive ratio of non-vaccinated pigs | 1 – 9.04 | [43, 49, 52, 55] |
| $R_0^V$ | Basic reproductive ratio of vaccinated pigs | 0 – 7.1 | Estimated by relations (3) & (7)–(9)<br>(see also Ref. [43, 49, 52]) |

The average number of secondary cases arising from one infection when the entire population is susceptible, i.e. the basic reproductive ratio, denoted by *R*_0_ is given by the following Eq [46]

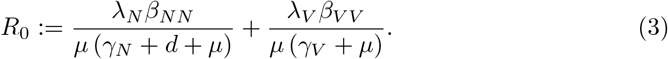

*R*_0_ is a key epidemiological measure for how “infectious” a disease is, with large *R*_0_ possibly representing more virulent pathogen strains [47]. Vaccination ultimately aims to prevent the invasion of an infection into a population, which corresponds to reducing *R*_0_ to below one (*R*_0_ < 1).

One of the factors that compromise vaccine effectiveness is the effective vaccine coverage, *p*, which may capture the proportion of herd members to which the vaccine has been applied or the proportion of herd members that respond to the vaccine, i.e. are effectively immunized (see Table 1). Vaccine coverage *p* is implemented into the model through the replacement rates.

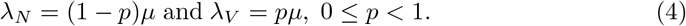

Therefore, for 0 ≤ *p* ≤ 1 the basic reproductive ratio in a population with effective vaccine coverage *p* becomes

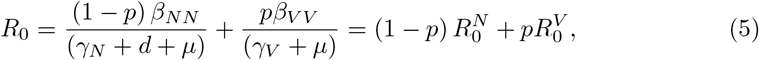

where 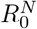 and 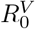 the basic reproductive ratios in non-vaccinated and vaccinated pigs, respectively. In the case of no removal or replacement, the basic reproductive ratio of the closed herd is

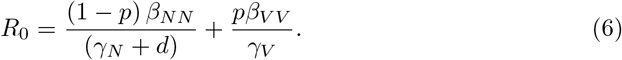

### Vaccine properties

To investigate vaccine effectiveness, we distinguish between different types of vaccines and different vaccination strategies (see Fig 2). In particular, we investigate how the diverse factors compromising vaccine effectiveness, in addition to effective vaccine coverage, listed in Table 1 separately and combined affect the infection invasion and spread in the herd. Vaccine properties either affect the transmission rates *β_jk_* or the recovery rates *γ_j_,j, k* = *N, V*, in the epidemiological model (1). Transmission rates may be reduced by the vaccine in two ways, i.e. either due to a reduction in host susceptibility, modelled by the vaccines sterilizing immunity *ϵ_s_*, or due to a reduction in host infectivity, modelled by a vaccine-induced effect *ϵ_i_* on the transmission rate. Note that these effects may refer to two different mechanisms. For example vaccine-induced reduction in host susceptibility may refer to mechanisms reducing pathogen entry and establishment into the target cells, whereas vaccine-induced reduction in host infectivity may refer to mechanisms regulating pathogen replication and shedding. Similarly, a vaccine may trigger immune mechanisms that speed up recovery. Thus, we also model here the vaccine-induced effect on the recovery rate denoted by *ϵ_γ_*.

**Fig 2.**
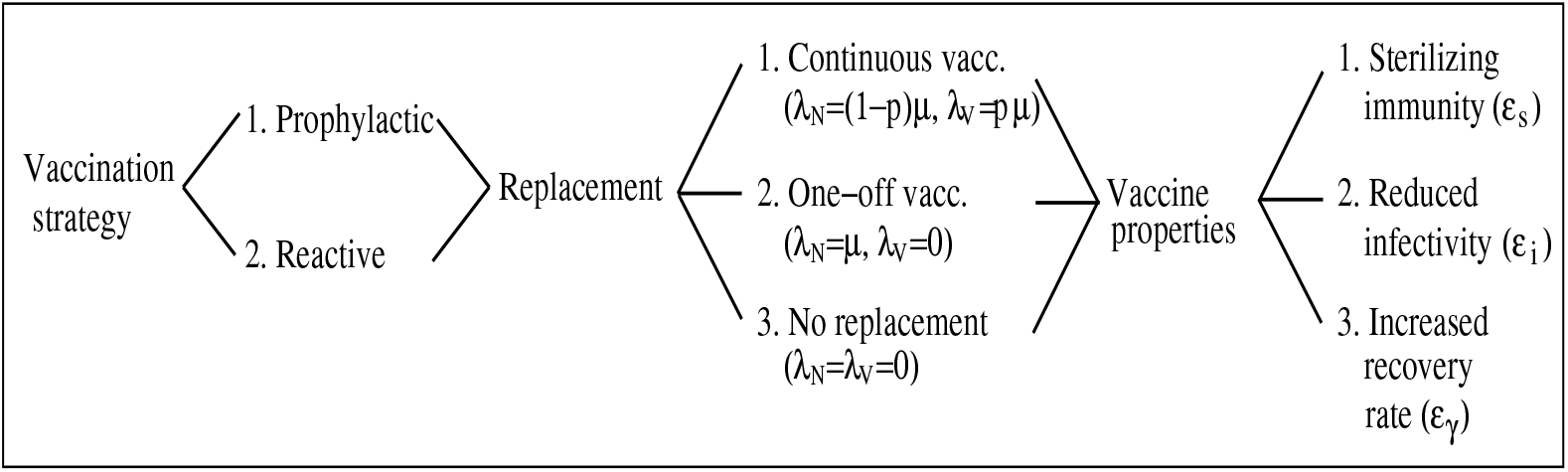
Tree diagram showing modelled scenarios. We refer to the text for definitions of the parameters and further explanations. Here, the vaccine coverage *p* represents either the proportion of individuals to which the vaccine is administered or the proportion of vaccine responders. Also, delay in the onset of vaccine induced immunity is not explicitly included as vaccine property as it is equivalent to modelling a reactive vaccination strategy.

Assuming that these vaccine properties act independently from each other (i.e. multiplicative effects on the model parameters), the different vaccine properties are represented in the model (1) as follows:

1. *Vaccine-induced sterilizing immunity ϵ_s_*(0 ≤ *ϵ_s_* ≤ 1): In this case the vaccine reduces the susceptibility of the vaccinated individual with the following effects on the transmission rates

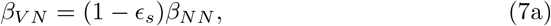

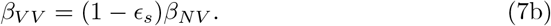
2. *Vaccine-induced reduction in host infectivity ϵ_i_*(0 ≤ *ϵ_i_* ≤ 1): In this case the vaccine reduces the propensity of a vaccinated infected host to transmit infection to a susceptible host upon contact, which is represented by

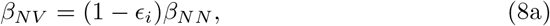

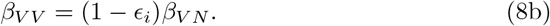
3. *Vaccine-induced increase in recovery rate ϵ_γ_* (0 < *ϵ_γ_* ≤ 1): This is modelled by

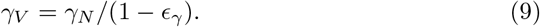
4. *Delay in the onset of vaccine-induced immunity:* The delay in the onset of vaccine-induced immunity is incorporated in the model by changing the constants *ϵ_s_, ϵ_i_* and *ϵ_γ_* to step functions *ϵ_s_*(*t*), *ϵ_i_*(*t*) and *ϵ_γ_*(*t*) with

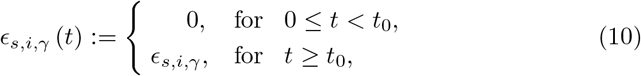

where *t*_0_ denotes the time period between vaccination and the onset of vaccine-induced immunity. As outlined below, these step functions are also used to model reactive vaccination. Finally, it should be noted that the limiting cases *γ_V_* → ∞ and *β_jk_* = 0 in equations (7)–(9) represent full protection of the vaccine and prevention of pathogen transmission when vaccination is applied, respectively.

### Vaccination strategies

In this study we distinguish between two types of vaccination strategies, i.e. prophylactic and reactive vaccination. In the case of prophylactic vaccination we assume that a proportion p of the herd has been effectively vaccinated at the time of exposure to the field pathogen strain. The initial conditions for this case are

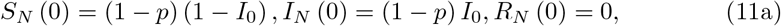

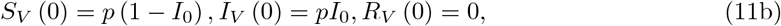

where *I*_0_ > 0 the total proportion of individuals (vaccinated and non-vaccinated) infected with the field pathogen strain. Thus in the case of full vaccine coverage *S_N_*(0) = *I_N_*(0) = *R_N_*(0) = 0.

Reactive vaccination refers to the situation when vaccination is applied only after a proportion of non-immunized individuals has become infected either due to delayed vaccination or a delayed onset of vaccine-induced immunization. It is modelled by the step functions (10), describing the effect of vaccination on transmission and recovery rates once the vaccination is applied at the time point *t*_0_ > 0.

Besides the two vaccination strategies, which mainly specify the proportion of vaccinated individuals in a specific period, another factor that controls transmission dynamics within a herd included in the model is the frequency of vaccination, and in particular whether vaccination is applied continuously or only once (hereafter denoted as one-off vaccination). In an open herd with constant replacement of removed or dead individuals, continuous versus one-off vaccination affect the replacement rates λ_*N*_ and λ_*V*_ as follows:

1. *Continuous vaccination in an open herd:* In this case we assume a continuous vaccination of incomers, with constant vaccine coverage *p*, i.e. λ_*N*_ = (1 − *p*)*μ*, λ_*V*_ = *pμ*.
2. *One-off vaccination in an open herd*: In this case we assume that vaccination is not applied to incoming animals, i.e. λ_*N*_ = *μ*, λ_*V*_ = 0.

In a closed herd, vaccination is assumed to be applied only once, and in this case the replacement rates λ_*N*_ and λ_*V*_ are zero, i.e. there are no incomers. In summary, different vaccination frequencies in open or closed herds can be modelled through different replacement rates.

Fig 2 illustrates the different scenarios corresponding to different vaccine properties and vaccination strategies modelled in this study.

### Application to PRRS

The model presented in equations (1)–(2) above can be applied to model PRRSv transmission dynamics in pig herds under different vaccination scenarios [48,49,51]. Values of the model parameters were obtained from the literature (see references listed in Table 2) and represent different PRRSv strains (e.g. low or highly virulent strains), different types of pig herds (e.g. sows vs growing pigs), and different management structures, etc. The large range in values for epidemiological parameters in non-vaccinated herds allows assessment of vaccine and vaccination effects under a wide range of conditions. We therefore present results for low and high *R*_0_ values *R*_0_ ≈ 2 and R_0_ ≈ 6, respectively.

In each simulated scenario, the infection in a herd is introduced by assuming 0.1% of the pigs are infected at the beginning of the observation period. Depending on the model scenario, these are either vaccinated (prophylactic vaccination) or non-vaccinated (reactive vaccination).

To assess vaccine effectiveness we focus on three important epidemiological measures [45] (i) the risk of infection invasion (or infection eradication if already invaded), represented by *R*_0_; (ii) the peak prevalence of infection; (iii) the time at which the peak prevalence occurs.

Table 2 lists the parameter value ranges used in the simulations together with the literature source. Here we show results for two different average values of 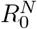, i.e. 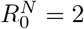 and 6 [43,49,52,55], representing a moderate and a severe PRRS epidemic, respectively. These values were produced by assuming an average duration of the infectious period of 56 days [49,55,56], corresponding to a recovery rate for non-vaccinated pigs of of *γ_N_* = 0.01785 per day. The assumed transmission rate corresponding to 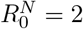 was *β_NN_* = 0.04 per day, whereas the transmission rate corresponding to 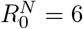 was *β_NN_* = 0.12 per day. Choosing different parameters sets of *β_NN_,γ_N_* can lead to the same 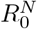 values, generalising a bit the results and giving a good theoretical framework, since presenting all the possible sets is not possible.

## Results

### 1 Prophylactic vaccination

#### 1.1 Continuous vaccination

To quantify the effect of a vaccine on infection invasion in a herd we use relation (5), which becomes

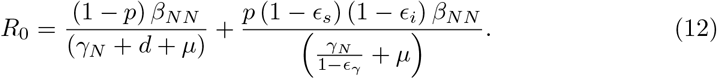

Assuming negligible infection induced death rate *d* (see Table 2), Eq (12) simplifies to:

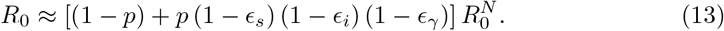

Relation (13) shows that the vaccine properties *ϵ_s_, ϵ_i_* and *ϵ_γ_* exhibit an equivalent effect on *R*_0_. Because of the assumed independence between these vaccine properties (Eq. (7)–(9)), different effects of a vaccine act multiplicatively on *R*_0_. Thus, assuming vaccine coverage of one (*p* = 1), a vaccine providing 50% sterilizing immunity (*ϵ_s_* = 0.5), but having no effect on the infectivity or recovery rate of individuals, reduces the R0 by approximately 50%. A vaccine that in addition to a 50% sterilizing immunity also reduces infectivity or speeds up recovery by 50%, leads to 75% reduction in *R*_0_, whereas vaccines that have a beneficial effect on all three host traits susceptibility, infectivity and recovery, reduces *R*_0_ by 88.5%. This symmetry is also illustrated in Fig 3A-B, which shows a surface plot (panel A) and a contour plot (panel B) of the dependence of *R*_0_ on vaccine-induced sterilizing immunity, *ϵ_s_*, and vaccine-induced effects on infectivity, *ϵ_i_*, and recovery, *ϵ_γ_*, respectively, for vaccine coverage *p* =1. The coloured surface/curves correspond to the threshold *R*_0_ = 1 in Eq (12), between scenarios in which the infection is expected to invade (*R*_0_ > 1; area below the threshold) or not invade (*R*_0_ ϵ 1; are above the threshold) a herd. The graphs show that even for large 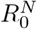 (e.g. virulent PRRSv strains), one of the investigated vaccine properties alone can prevent the infection invasion if the effect size is sufficiently large (e.g. *ϵ_s,i_* ≥ 0.84 or *ϵ_γ_* > 0.85 for 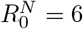). However, considerably less sterilizing immunity or reduction in infectivity are required, if the vaccine also simultaneously speeds up recovery (e.g. *ϵ_s,i_* ≥ 0.5 is sufficient if *ϵ_γ_* = 0.7).

**Fig 3.**

The dependence of *R*_0_ on the vaccine-induced sterilizing immunity, *ϵ_s_*, vaccine-induced reduction in host infectivity, *ϵ_i_*, vaccine-induced increase in recovery rate, *ϵ_γ_*, and vaccine coverage, *p*. A. 3D surface plot corresponding to *R*_0_ = 1 in Eq (12), for a high average transmission potential 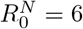 and full immunization coverage (*p* = 1). B. 2D contour plot showing the dependence of *R*_0_ on *ϵ_i_* and *ϵ_γ_*, for different values of *ϵ_s_* and *p* =1. C. 3D surface plot corresponding to *R*_0_ = 1 in Eq (12), for a high average transmission potential 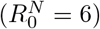 and *ϵ_i_* = 0.5. D. 2D contour plot showing the dependence of *R*_0_ on the effective vaccine coverage, *p*, and *ϵ_γ_*, for different values of *ϵ_s_* and *ϵ_i_* = 0.5. The coloured continuous curves in 2D plots correspond to *R*_0_ = 1 in Eq (12) for a high average transmission potential 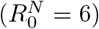 and dotted curves correspond to *R*_0_ = 1 for a low virulent PRRSv strain 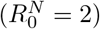. Areas above these curves correspond to *R*_0_ < 1 where the infection cannot invade the herd. Here *d* = 0.001, *μ* = 0.0017. The transmission and recovery rates are values within the ranges given in Table 2, that satisfy 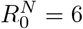 and 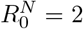 for the continuous and dotted curves, respectively.

Fig 3C-D show equivalent plots for incomplete vaccine coverage. The graphs demonstrate that infection invasion can be prevented even if the vaccine offers low or zero vaccine-induced sterilizing immunity, given sufficiently high vaccine coverage and vaccine-induced effect on host infectivity or recovery. For the plots in Fig 3C-D it was assumed that *ϵ_i_* = 0.5, i.e. the vaccine reduces host infectivity by 50%. For lower values of *ϵ_i_* higher levels of effective coverage or/and vaccine-induced increase in recovery rate would be needed to prevent infection invasion for given levels of sterilizing immunity. Only when the vaccine is fully protective (i.e. any of *ϵ_s,i,γ_* = 1) the required vaccine effective coverage no longer depends on the other vaccine effects, but only on the 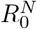 value in the equivalent non-vaccinated population (see gold continuous and dotted lines in Fig 3B). For example, for 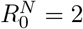, vaccination with a fully protective vaccine would require at least 49% vaccine coverage, whereas 83% coverage would be required for 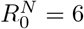 (Fig 3D).

In contrast to the near symmetric effect of the different vaccine properties, represented by *ϵ_s,i,γ_*, on *R*_0_, their effect on the infection dynamics can be diverse. Both vaccine-induced sterilizing immunity and vaccine-induced reduction in host infectivity have a symmetric effect on the transmission rate (Eqs (7) and (8)). However, Fig 4 shows that the effects of *ϵ_s_* and *ϵ_i_*, on the infection dynamics are different to those corresponding to *ϵ_γ_*. Higher vaccine-induced sterilizing immunity, *ϵ_s_*, or greater reduction in infectivity, *ϵ_i_*, (not shown here) leads to milder, but prolonged epidemics due to the slower rate at which infection is transmitted, causing also a later occurrence of the first infection incidence and peak prevalence, and a slower rate of post-peak prevalence decline (Fig 4A). In contrast, vaccine-induced increase in recovery rate, *ϵ_γ_*, mainly affects peak prevalence, but not the timing at which it occurs or the rate of decline (Fig 4C). For continuous prophylactic vaccination, vaccine-induced effects on transmission or recovery rates also affect the long-term steady-state prevalence of infection. Higher vaccine-induced effect on transmission or recovery rate lead to a greater reduction of the long-term endemic steady state. For example a 50% vaccine-induced sterilizing immunity, *ϵ_s_* (or 50% vaccine-induced reduction in host infectivity, *ϵ_i_*), will reduce the endemic steady state of the infected animals by 25%, while a 50% vaccine-induced increase in recovery rate will reduce the endemic steady state of the infected animals by 59%. As would be expected, multiple vaccine properties combined affect both the timing and level of peak infection prevalence, and the rate of decline (Fig 4E). The infection dynamics also confirm the results shown in Fig 3A-B, that an 85% of susceptibility, infectivity or recovery alone (Fig 4A,C) or a 50% combined effect of all (Fig 4E) is required to prevent infection from invading a herd.

**Fig 4.**
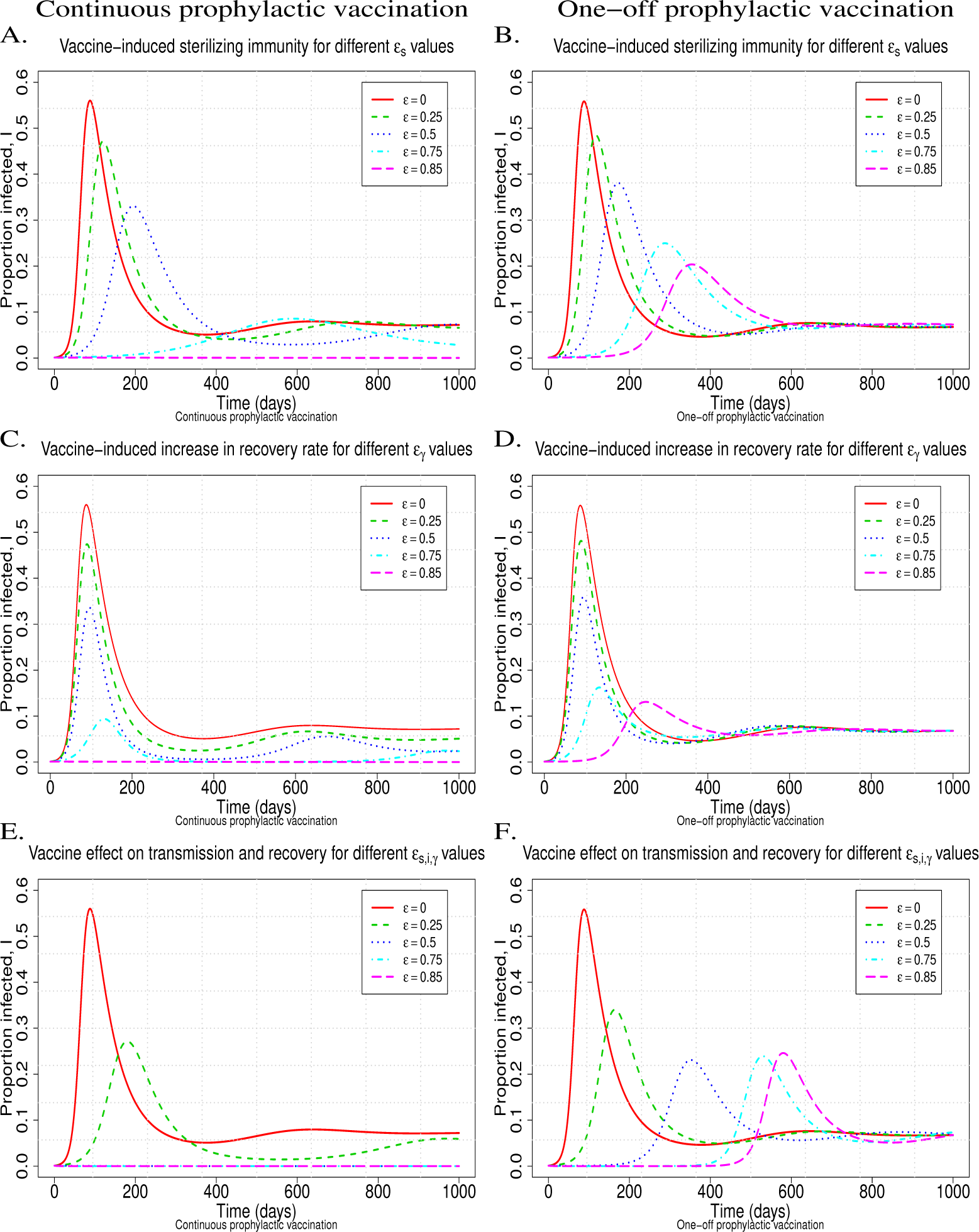
The effect of prophylactic mass vaccination for different *ϵ* = *ϵ_s,iγ_* values on the infection dynamics for continuous vaccination, i.e. λ_*N*_ = 0, λ_*V*_ = 0.0017 (left column), and one-off vaccination, i.e. λ_*N*_ = 0.0017, λ_*V*_ = 0 (right column). A.-B. The vaccine only affects host susceptibility, i.e. sterilizing immunity *ϵ* = *ϵ_s_, ϵ_γ_* = *ϵ_i_* =0 (see relation (7)); C.-D. The vaccine only affects recovery rate, i.e. *ϵ* = *ϵ_γ_, ϵ_s_* = *ϵ_i_* = 0 (see relation (9)); E.-F. The vaccine equally affects host susceptibility, infectivity and recovery rate, *ϵ* = *ϵ_s,i,γ_* (see relations (7)-(9)). Other chosen parameter values were *β_NN_* = 0.12, *γ_N_* = 0.01785, *d* = 0.001, *μ* = 0.0017, and the initial conditions are *S_V_* (0) = 0.999, *I_V_* (0) = 0.001 (see relation (11) for *p* =1 and *I*_0_ = 0.001).

#### 1.2 One-off vaccination

One-off vaccination in an open herd with constant replacement rate implies that λ_*V*_ =0 and thus does not reduce the basic reproductive ratio, i.e. 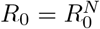 (see relation (3)). This is due to the immediate influx of non-vaccinated pigs into the herd. If there is a continuous influx of non-vaccinated pigs into the herd, one-off vaccination is not adequate for preventing infection invasion. However, as shown in Fig 4 (right column), vaccination can substantially reduce the severity of the epidemic. For example, in the simulations of Fig 4, a 50% vaccine effect on susceptibility, infectivity and recovery can reduce the peak prevalence up to 57.1% (Fig 4F). Moreover, the greater the effect of a vaccine on host susceptibility or infectivity, the later the occurrence of peak prevalence (Fig 4B). Comparison of the continuous (left-column) with the one-off (right-column) prophylactic vaccination shows that the short-term effects are very similar in both cases (Fig 4). In contrast, whereas in the one-off vaccination scenario, the endemic steady state is not affected by vaccination (i.e. the endemic steady state is the same for every value of 0 ≤ *ϵ_s,i,γ_* < 1; Fig 4B,D,F), continuous vaccination alters the endemic steady state values, whereby more effective vaccines (i.e. greater values of *ϵ_s,i,γ_*; 4A,C,E) correspond to lower prevalence in the long-term.

#### 1.3 Closed herd

Based on relation (6), the effect of vaccination on infection invasion (or eradication if already invaded) in a closed herd is given by

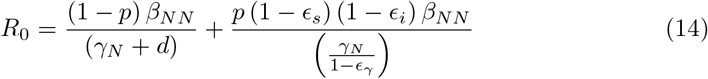

which is the same as *R*_0_ for the continuous vaccination of a herd with replacement when *μ* = 0 (see Eq (12)). Therefore, the vaccine properties, *ϵ_s,i,γ_*, will have a similar effect on preventing an infection to invade a herd as the continuous vaccination in an open herd. This can be easily seen by comparing the left columns of Figs 4 and 5 (prophylactic vaccination in a closed herd). In other words, even for severe epidemics in non-vaccinated herds with corresponding 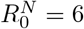, a vaccine with sufficiently strong effect on either host susceptibility, infectivity or recovery can prevent infection invasion (e.g. *ϵ_s,i,γ_* ≥ 0.852 for 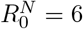), and this can also be achieved by a combination of effects with weaker effect size (e.g. *ϵ_s_* = *ϵ_i_* = *ϵ_γ_* = 0.5 for 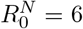). The main difference of a closed herd compared to an open herd is that in a closed herd, the *R*_0_ value corresponding to the same values for the epidemiological model parameters (*β_jk_, γ_j_, j, k* = *N, V* and *d*) is greater and thus the infection invasion is more likely. This implies that for a closed herd, vaccine requirements for preventing the invasion of infections are stricter than for open herds. Differences in the infection dynamics between open and closed herds occur mainly in the long term, where the infection dies out in closed herds, regardless of whether or not vaccination is applied, whereas it reaches an endemic equilibrium in open herds (Fig 4 and Fig 5 and [17]).

**Fig 5.**
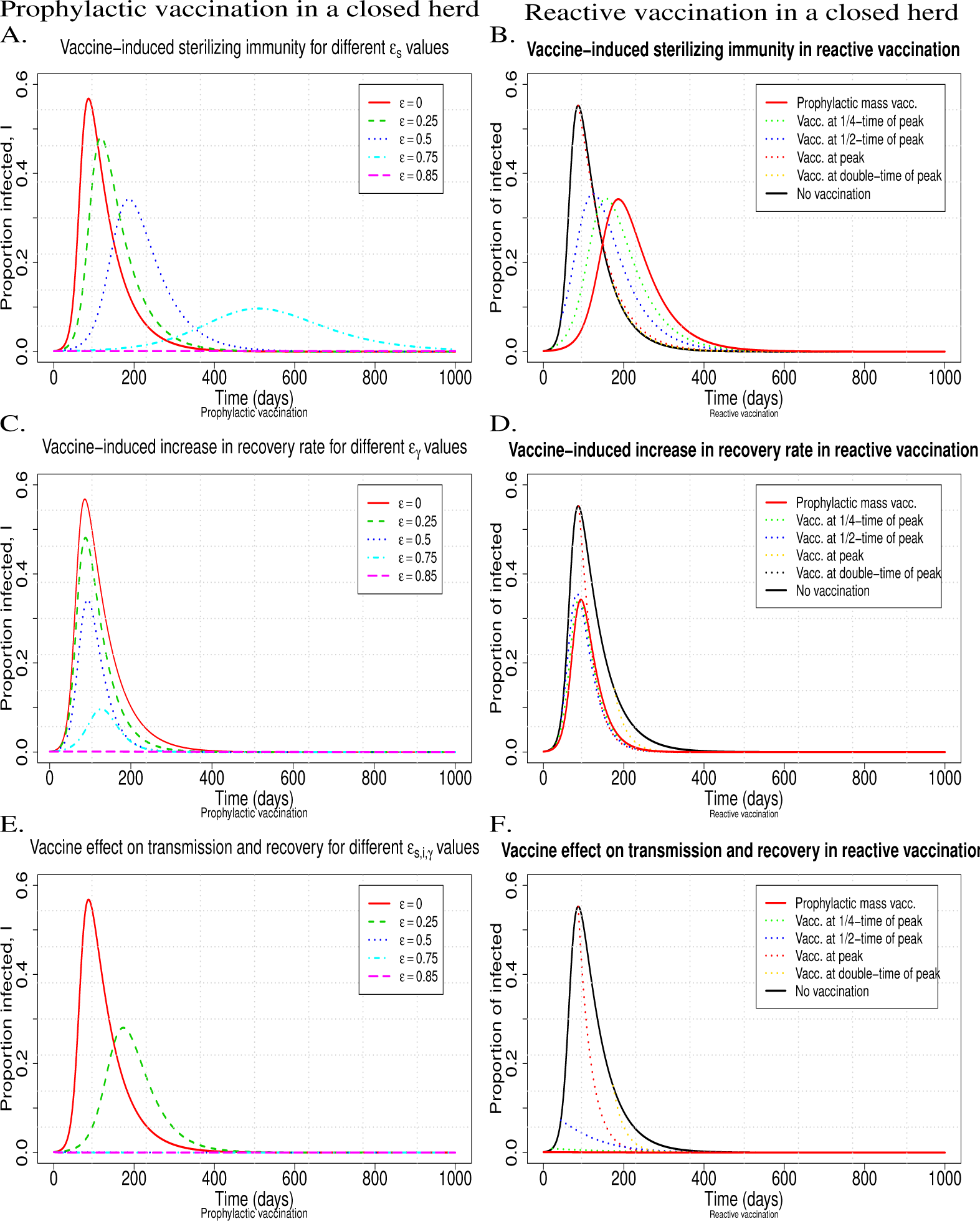
The effect of fully effective mass vaccination (i.e. *p* = 1 at the time of vaccination) with vaccines of different properties on the infection dynamics in the case of prophylactic (left column) and reactive (right column) vaccination in a closed herd. Profiles of different colours correspond to different *ϵ* = *ϵ_s,i,γ_* values on the infection dynamics (left column; see caption in Fig 4), or different times when vaccination is applied (right column; see caption in Fig 6). A.-B. The vaccine only affects host susceptibility, i.e. sterilizing immunity, but no effect on infectivity or recovery (see Eq (7) and (10)); C.-D. The vaccine only speeds up recovery, but offers no protection from infection and no reduction in infectivity (see Eq (9) and (10)); E.-F. The vaccine equally affects host susceptibility, infectivity and recovery rate (see Eq (7)–(9) and (10)). Here β_*NN*_ = 0.12, γ_*N*_ = 0.01785, *p* = 1, *d* = 0.001, *μ* = λ_*N,V*_ = 0, and the initial conditions are *S_V_* (0) = 0.999, *I_V_* (0) = 0.001 (left column) and *S_N_* (0) = 0.999, *I_N_* (0) = 0.001 (right column).

### 2 Reactive Vaccination

In modelling terms, reactive vaccination is equivalent to applying prophylactic vaccination with delayed onset of immunity at a stage where infection invades a herd before the full vaccine-induced immunization effects have been reached. Thus, although these types of vaccination cannot prevent the infection from invading, they can still reduce the prevalence of infection and may ultimately eliminate the infection from a herd. Fig 6 shows prevalence profiles resulting from reactive vaccination in a herd with replacement, either applied continuously (left column) or one-off (right column). Overall, the vaccine effects on the infection dynamics are similar to those observed for the corresponding prophylactic vaccinations (see Fig 4), with continuous vaccination leading to a long-term reduction in the prevalence of infection and one-off vaccination only affecting infection prevalence in the short-term, but reaching the same endemic equilibrium as that corresponding to the equivalent non-vaccinated herd (Fig 6). Reactive vaccination, even if applied as a one-off disease control, can substantially reduce peak prevalence. In fact, prevalence profiles corresponding to reactive vaccination applied prior to the time of peak prevalence, look remarkably similar to those corresponding to prophylactic vaccination with the same vaccine, although prevalence generally reaches its peak slightly later in the prophylactic vaccination scenario (Fig 6A-F). Moreover, reactive vaccination, when applied continuously can eliminate infection in a herd, under the same required vaccine properties as for prophylactic vaccination, regardless of when it is applied (Fig 6E). For example, Fig 6E shows a scenario where the infection can be eliminated from a herd when reactive vaccination inducing 50% combined vaccine effect on transmission and recovery rate is applied. The timing of the reactive vaccination affects primarily peak prevalence and total number of infectees (early vaccination corresponds to lower the peak prevalence and less infectees), rather than whether and when the infection is eliminated from the herd (Fig 6).

**Fig 6.**
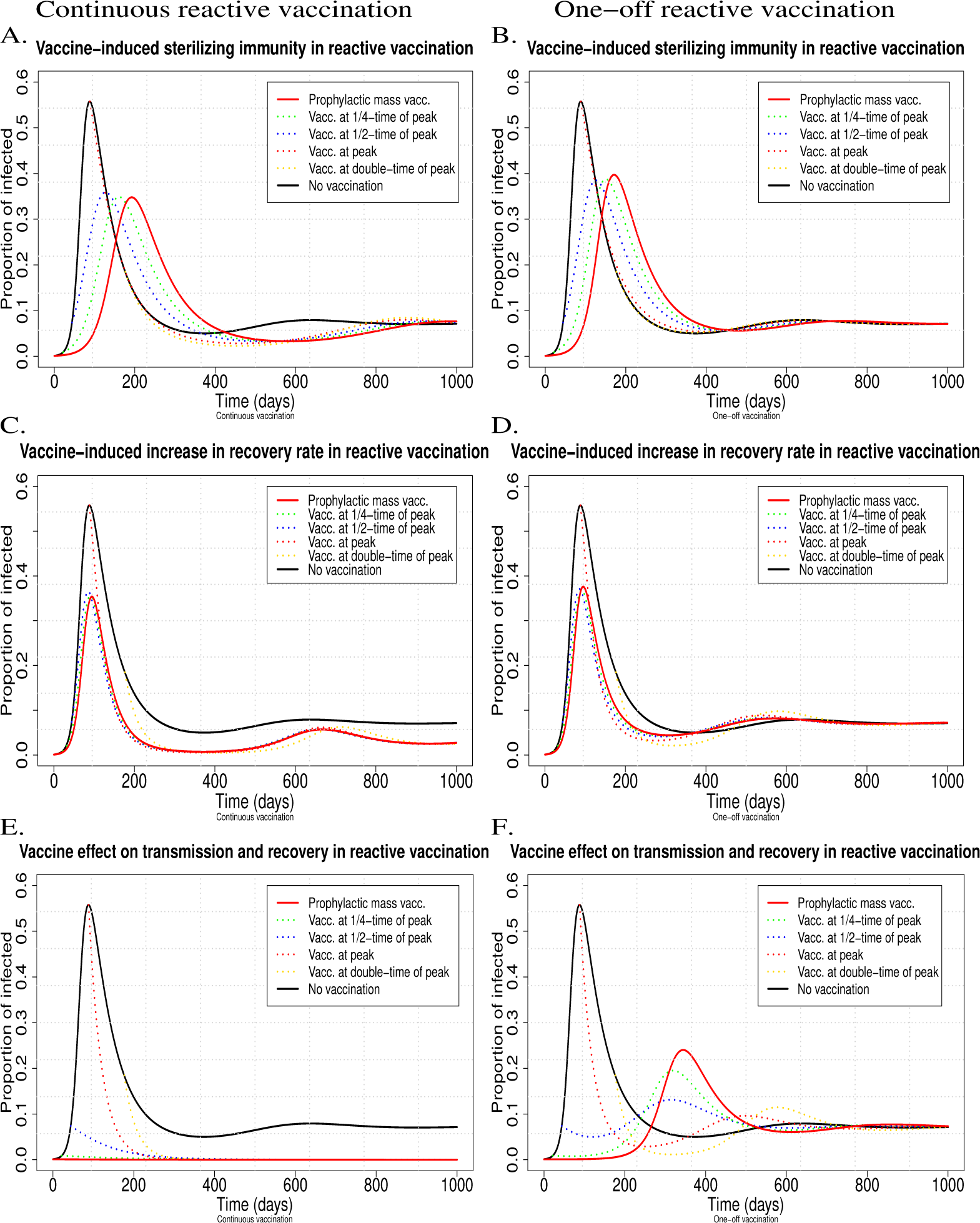
The effect of complete mass vaccination (i.e. *p* = 1) with vaccines of different properties on the infection dynamics in the case of continuous (left column) and one-off (right column) reactive vaccination. Profiles of different colours correspond to different times when vaccination is applied, i.e. (i) at the start of the epidemic, equivalent to prophylactic vaccination (red continuous curves), (ii) at one quarter of the time of peak prevalence (green dotted curves), (iii) at one half of the time of peak prevalence (blue dotted curves), (iv) at the time of peak prevalence (red dotted curves), (v) at double the time until peak prevalence (gold dotted curves) and when (vi) vaccination is not applied at all (black continuous curves). A.-B. The vaccine only affects host susceptibility, i.e. sterilizing immunity, but no effect on infectivity or recovery (*ϵ_s_* = 0.5, *ϵ_γ_* = *ϵ_i_* = 0) (see relations (7) and (10)); C.-D. The vaccine only speeds up recovery, but offers no protection from infection and no reduction in infectivity (*ϵ_γ_* = 0.5, *ϵ_s_* = *ϵ_i_* = 0) (see Eq (9) and (10)); E.-F. The vaccine equally affects host susceptibility, infectivity and recovery rate, *ϵ* = *ϵ_s,i,γ_* = 0.5 (see Eq (7)–(9) and (10)). Here *β_NN_* = 0.12, *γ_N_* = 0.01785, *p* =1, *d* = 0.001, *μ* = 0.0017, and the initial conditions are *S_N_* (0) = 0.999, *I_N_*(0) = 0.001 (see relation (11) for *p* = 0 and *I*_0_ = 0.001).

Somewhat surprisingly, Figs 6B and 6F indicate a slightly higher peak prevalence in the case of one-off prophylactic mass vaccination compared to one-off reactive mass vaccination when applied early in the epidemics. This is because the time and level of peak prevalence partly depends on the relative proportion of infected animals when the vaccination is applied and the proportion of new non-vaccinated incomers. At the time when prophylactic vaccination is applied the initial proportion of infected vaccinated animals is very low and infection can only spread slowly initially. However, prevalence rises relatively quickly as the non-vaccinated replacement become infected and start to transmit the infection. In contrast, at the time when reactive vaccination is applied (e.g. at 1/4 of peak prevalence time) all infected non-vaccinated animals become vaccinated with lower transmission rate (assuming *ϵ_i_* > 0), so that the proportion of new non-vaccinated incomers will not face many highly infectious animals in the herd. Thus, this case illustrates that a one-off reactive vaccination applied at the right time of an ongoing epidemic can be more effective than the one-off prophylactic vaccination.

Fig 5 shows that in a closed herd the vaccine properties, *ϵ_s,i,γ_*, will have a similar effect on eradicating an infection as the continuous reactive vaccination in an open herd (see left column in Fig 6). Similar to the prophylactic vaccination, the main difference of a closed herd to an open herd is that the infection will eventually die out in all vaccination scenarios (even without vaccination) in a closed herd.

## Discussion and Conclusion

Imperfect vaccines that do not prevent infection transmission and heterogeneous response to vaccines are common in livestock production and threaten the success of vaccination programs [57–59]. It is well known that the success of vaccination programs depends on the effective vaccination coverage as well as on the type of vaccine (e.g. [60]). However, vaccination programs differ not only in these components, but also in when and how a particular vaccine is applied as a single control measure or in combination with others such as herd closure. Yet, to date relatively little is known how these factors interact with each other and affect the outcome of vaccination programs. The epidemiological model developed in this study provides for the first time detailed quantitative understanding of the relationship between these diverse factors and disease invasion and spread in domestic livestock. The model predicts within-herd infection dynamics in livestock populations following the application of imperfect vaccines, and provides threshold conditions for vaccine properties and vaccination strategies to prevent infection invasion or eliminate infection from a herd after infection invasion. This model was parameterised and applied to PRRS as one of the major livestock examples where imperfect vaccines are commonly applied and much debated in the scientific literature [8,25,26].

Most previous modelling studies have mainly focused on differences in vaccine efficacy captured by reduction in *β*. Vaccine efficacy is however multi-faceted and most vaccines reduce pathogen transmission not only by offering sterling immunity, but also by reducing host infectivity and duration of infection. We therefore distinguished between multiple effects in our model and we found that a vaccine can deviate substantially from perfect in one or all of these properties, as long as they reduce pathogen transmission in several ways. For instance for disease with an average transmission potential of *R*_0_ = 6 in a non-vaccinated herd, we found that vaccines that reduce susceptibility, infectivity and recovery rate by 50% could prevent infection invasion or eradication invaded infection. Moreover, our model revealed that imperfect vaccines, that only offer partial protection from infection with a heterologous virus strain, can substantially reduce pathogen transmission and infection risk and prevalence in herds, when applied as prophylactic strategy. Unsurprisingly, our model predicts that prophylactic mass vaccination with a vaccine that confers high sterilizing immunity in all individuals is the most effective strategy for preventing infection in a herd or for reducing prevalence. However, our model also shows that even vaccines that offer no or little sterilizing immunity or only provide partial effective coverage can prevent infection invasion into a herd if the vaccine simultaneously reduces host infectivity e.g. by reducing pathogen shedding or speeding up recovery time. In this case the occurrence of secondary infection cases can be successfully prevented, as indicated by a reduction of the basic reproductive ratio of the vaccinated herd to values below 1. The results are relevant to certain MLV vaccines showing low sterilizing immunity [8,21,26,31,32], but significant reduction in viral shedding and infectious period. The model results suggest that these vaccines, despite far from perfect, could achieve drastic reduction in the occurrence and severity of PRRS outbreaks in commercial pig populations, and even help to eliminate the disease.

As herd closure is a common control measure, a vaccination strategy considering a closed herd was studied. According to our model results, similar levels of reduction in infection peak prevalence can be achieved with one-off vaccination in closed herds as with continuous vaccination of herds with constant replacement rate, for the same type of vaccine. However, when prophylactic vaccination is applied in herds with constant influx of new animals, infection prevention can only be achieved through continuous vaccination. One-off vaccination may be used to delay and reduce peak prevalence, but cannot reduce long-term prevalence and hence eliminate the infection from the herd.

Both, prophylactic and reactive vaccination are routinely applied to control PRRS in pig farms [25,61]. Reactive vaccination in particular is commonly used in farms where the virus has been found endemic [8, 26]. Our model results indicate that the timing of vaccination is crucial for achieving effective reduction in the prevalence of infection, whereas the frequency of vaccination controls the chance of eliminating the infection from a herd. Indeed, reactive vaccination campaigns are racing against the timeline in which the infection spreads within a herd, and timely distribution of vaccines in response to an ongoing epidemic can prevent new infection cases [20,62,63]. In our model, continuous reactive vaccination with a vaccine that has combined positive effects on host transmission and recovery rates could eliminate the infection from a herd even for high average transmission potential characterized by high 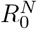 in non-vaccinated populations. As PRRS KV vaccines are known to elicit immune responses to the infecting virus in PRRSv-positive pigs (reactive vaccination) [40,41], and offer low sterilizing immunity in PRRSv-negative pigs, they are often not considered as effective vaccines for disease prevention (i.e. PRRS KV vaccines are no longer available in the United States since 2005 [8]), but might exert a potential role as a therapeutic vaccine for PRRSV treatment [8,26]. However, immune studies have shown further that KV vaccines can reduce pathogen shedding in seropositive animals, hence host infectivity, and speed up recovery [42]. In line with this study, our model suggests that PRRS KV vaccines can still effectively reduce the transmission of the infection or eliminate infection from herds by reducing pathogen shedding and speeding up recovery. Thus, from a purely epidemiological perspective, the results would imply that KV, if adequately applied could lead to drastic reduction infection prevalence. As prophylactic vaccination is often avoided in practice in order to reduce the risk of side effects from vaccination (e.g. vaccination alone may cause low percentages of abortions when used in breeding herds), timely and targeted reactive vaccination may be the most effective way to mitigate PRRS transmission within herds. The large differences in vaccine effectiveness associated with different vaccination strategies indicate the importance of designing vaccination programs appropriately as vaccine effectiveness depends largely on the timing and frequency of vaccination, in addition to the particular vaccines properties.

The model developed in this study is a generic epidemiological SIR model adapted to PRRS by adopting a large range for the model parameter values from the literature to represent PRRSv strains of different virulence, different pig populations and different management structures. This generic modelling approach combined with large parameter ranges was deliberately chosen to gain relevant qualitative insight of vaccine and vaccination effects for a wide scope of scenarios. Obtaining reliable quantitative predictions for specific situations would require obtaining accurate estimates of the epidemiological model parameters *β* and *γ* and the vaccine effects *ϵ_s_, ϵ_i_* and *ϵ_γ_* for existing vaccines; although inferring these from existing data may be challenging and may require specific experimental designs [43,52,55]. Other model limitations include that specific herd or farm structure, known to influence within-herd transmission dynamics of pathogens (see e.g. [48,49,51]), were not explicitly considered in our model. Furthermore, we only considered horizontal infection transmission in this study, thus ignoring the potential impact of vaccination on vertical transmission from infected sows or boars to piglets from the time of insemination during gestation until after birth [25,64]. One of the main aims of PRRSv vaccines is to reduce the impact of infection on growth performance in growing piglets or reproductive failure in pregnant sows [21,65]. Like most epidemiological models, our model does not explicitly distinguish between infection and disease, and hence additional parameters would be required to model the relationship between disease prevalence and productive or reproductive herd performance. Finally, the model developed in this study considered only transmission of a single wild-type strain in vaccinated populations, but PRRSv is a diverse quasi-species and different PRRSv strains can circulate simultaneously in single a herd [66,67]. The effects of partial protection against single strains are considered in this model, but the effects of heterogeneous protection against multiple variants simultaneously are not included in the present study. This may affect virus evolution and vaccine safety, and therefore also the predicted long-term effects of vaccination.

In the past, several mathematical models studied the transmission of PRRS in a homogeneous population [49,55,56], while other studies [43,48,51] focused on populations where vaccination is applied (considered heterogeneous in the sense of vaccination). These studies provide insight on the evaluation of transmission and recovery from PRRSv infection, and on the effect of vaccination on epidemic risk and severity. Many of them have a more explicit representation of the farm structure with detailed description of the demographics. Building on simple compartmental models, we performed a comprehensive study of different key factors that compromise the effectiveness of an incomplete vaccine, where our focus was to investigate the interactive effects of vaccine properties and vaccination strategies rather than modelling specific demographic properties. However, the high sensitivity or our model results on the replacement rate parameters indicate that demographic properties would need to be adequately represented to achieve high predictive power.

In summary, this study presents a systematic generic modelling framework to investigate the effectiveness of imperfect vaccines for preventing, mitigating or eliminating infectious diseases in animal populations as a function of vaccine properties, vaccination strategies and replacement rate of vaccinated and non-vaccinated individuals. The model results suggest that even imperfect vaccines with no or low levels of sterilizing immunity, or less than 100% effective coverage, when appropriately applied can prevent, eliminate or largely reduce the prevalence of PRRSv and other virus infections, as long as the vaccine sufficiently speeds up recovery and reduces pathogen shedding. The nearly multiplicative effect of diverse vaccine properties on the *R*_0_ in continuous vaccination highlights the importance of considering the combined effects of diverse vaccine properties in preventing infection invasion if applied prophylactically, or in eradicating the infection if applied reactively. In contrast, one-off vaccination with incomplete vaccines only cause a limited short-term reduction in infection prevalence, in particular in populations with high replacement rates. Overall, although continuous prophylactic mass vaccination is the most effective strategy in preventing infection invasion, one-off reactive vaccination can be more effective when applied at the right time of an ongoing epidemic than applied one-off prophylactically. The results have practical implications for the design of vaccines and vaccination programs in livestock populations. In particular, they suggest that in the absence of evolutionary constraints, the control or even elimination of PRRS through vaccination may well be within reach.

## Supporting information

**S1 Interactive modelling app** An interactive app for the model dynamics in the prophylactic vaccination under different replacement rates and different vaccine properties is available as an R Shiny app from (link).

## Acknowledgments

Financial support for this research was provided by the EU Horizon 2020 project SAPHIR, Project No. 633184 (VB and ADW), by the BBSRC Institute Strategic Programme Grants ISPG 2, Theme 2 (no. BBS/E/D/20002173) (SL, TO and ADW) and by the University of Edinburgh Chancellors Fellowship (SL). We would like to thank the coordinator of the SAPHIR project Prof. Isabelle Schwartz for her constructive comments to the manuscript.

